# Functional immune mapping with deep-learning enabled phenomics applied to immunomodulatory and COVID-19 drug discovery

**DOI:** 10.1101/2020.08.02.233064

**Authors:** Michael F. Cuccarese, Berton A. Earnshaw, Katie Heiser, Ben Fogelson, Chadwick T. Davis, Peter F. McLean, Hannah B. Gordon, Kathleen-Rose Skelly, Fiona L. Weathersby, Vlad Rodic, Ian K. Quigley, Elissa D. Pastuzyn, Brandon M. Mendivil, Nathan H. Lazar, Carl A. Brooks, Joseph Carpenter, Brandon L. Probst, Pamela Jacobson, Seth W. Glazier, Jes Ford, James D. Jensen, Nicholas D. Campbell, Michael A. Statnick, Adeline S. Low, Kirk R. Thomas, Anne E. Carpenter, Sharath S. Hegde, Ronald W. Alfa, Mason L. Victors, Imran S. Haque, Yolanda T. Chong, Christopher C. Gibson

## Abstract

Development of accurate disease models and discovery of immune-modulating drugs is challenged by the immune system’s highly interconnected and context-dependent nature. Here we apply deep-learning-driven analysis of cellular morphology to develop a scalable “phenomics” platform and demonstrate its ability to identify dose-dependent, high-dimensional relationships among and between immunomodulators, toxins, pathogens, genetic perturbations, and small and large molecules at scale. High-throughput screening on this platform demonstrates rapid identification and triage of hits for TGF-β- and TNF-α-driven phenotypes. We deploy the platform to develop phenotypic models of active SARS-CoV-2 infection and of COVID-19-associated cytokine storm, surfacing compounds with demonstrated clinical benefit and identifying several new candidates for drug repurposing. The presented library of images, deep learning features, and compound screening data from immune profiling and COVID-19 screens serves as a deep resource for immune biology and cellular-model drug discovery with immediate impact on the COVID-19 pandemic.

## INTRODUCTION

Acting through autocrine, paracrine, and endocrine mechanisms, endogenous immune stimuli maintain homeostasis and signal response to invasion, injury, or malignancy. Immune dysregulation underlies a broad set of human diseases including inflammation^1^, autoimmune disease^2^, neuroinflammation^3^, neurodegenerative disease^4^, secondary effects of traumatic brain injury^5^, cancer^6,7^, infection^8–10^, and cytokine storm^11,12^. Improvements in the understanding of how immune stimuli amplify or suppress the immune system, trigger new cell fate differentiation, and remodel tissue have resulted in the discovery of a wide range of successful therapeutics^13^, as demonstrated by the anti-TNF antibody adalimumab (Humira), noted both for its discovery^14^ and its application in rheumatic disease^15^. However, the immune system is vastly complex and dependent on cell type and context; reliably intervening in such a highly interdependent process is a formidable drug discovery challenge.

With few exceptions^16^, intercellular immune signaling has been explored by studying specific factors in isolation. Although cellular response to individual immune stimuli can be effectively profiled by high-dimensional molecular methods such as RNAseq^17^, proteomics^18^, and CHIPseq^19^, these technologies lack the speed and cost-effectiveness for systems-level analysis of large panels of immune stimuli and screening libraries across myriad inflammatory models. By contrast, image-based methods have demonstrated utility at many stages of drug discovery^20^, including compound profiling^21^, prediction of assay performance^22^, clustering by mechanism of action^23,24^, and toxicology predictions^25^.

Here we present phenomics, the analysis of fluorescence microscopy images as a scalable approach to examine cellular response to a wide range of perturbations. Deep-learning algorithms extract high-dimensional and dose-dependent fingerprints of cellular morphological changes, or ‘phenoprints’, from images to support a variety of downstream applications. These phenoprints detect subtle morphological changes far beyond human ability, and a standardized assay pipeline allows the phenoprints of millions of cellular samples to be related across time and experimental conditions. In this work, we first demonstrate the ability to use images alone to accurately quantify and relate hundreds of immune stimuli. We then show how profiles resulting from these perturbations can be employed in drug screening, particularly highlighting the utility of a relatable dataset to accurately predict the mechanism of action for unknown compounds. We used these capabilities to rapidly develop high-throughput-ready disease models for both SARS-CoV-2 viral infection and the resulting cytokine storm, and immediately launched large-scale drug screens that recapitulated known effective and ineffective therapies and, more importantly, identified several new potential treatments for both SARS-CoV-2 infection and COVID-19-associated cytokine storm.

## RESULTS

### Mapping response to immune stimuli: Generating phenoprints

In order to discern the breadth of immune response that might be captured in a single profiling assay for various applications (Fig. 1), we treated endothelial cells, fibroblasts, peripheral blood mononuclear cells (PBMC), and macrophages with diverse immune stimuli (Table S1), CRISPR gene editing reagents, antibodies, and small molecules. We used a single fully automated experimental workflow in which cells are plated, treated, labeled with a panel of fluorescent stains, and imaged with high-throughput fluorescence microscopy^26^. Vector representations of more than one million multi-channel fluorescence microscopy images analyzed in this manuscript were generated using a proprietary analytics workflow based on an extension of a DenseNet-161 neural network^27^

**Fig. 1:**
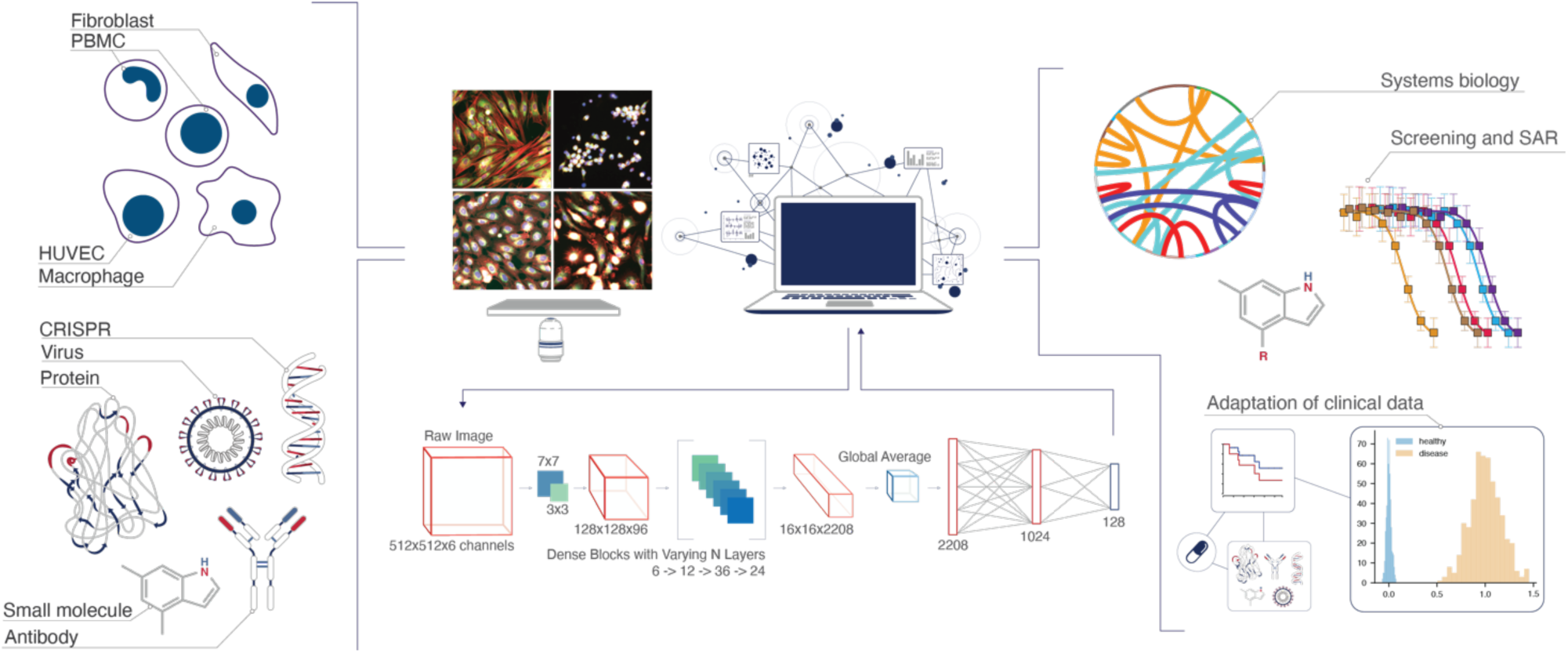
Phenomic platform for scaled discovery and exploration in immunology. Various cell types (top left) are treated with a range of biological perturbants and treatments (bottom left), including recombinant proteins, antibodies, CRISPR-based genetic modifications, and small molecules. High-throughput fluorescence microscopy (middle-top) and deep learning-enabled image featurization generates high-dimensional phenoprints that are used for interrogating a range of experimental questions (middle-top and middle bottom). This approach suits a suite of applications (right) with a single workflow and on a single platform. Specific applications demonstrated in this paper: systems biology exploration of immune relationships, adaptation across diverse disease models, compound screening, and mechanism prediction.

### Identifying immune stimulant phenoprints

Phenoprints are high-dimensional vector representations (embeddings) of cellular morphology derived from fluorescent images resulting from treatment with a biological perturbation. To provide landmark phenoprints across a diverse range of immune function, 446 stimuli were added to four different primary human cell types at a range of biologically relevant concentrations and compared to untreated cells and adjacent concentrations. For example, cells treated with tumor necrosis factor alpha (TNF-α), the anti-PD-1 antibody nivolumab, transforming growth factor beta (TGF-β), or interferon alpha (IFN-α) each displayed concentration-dependent phenotype strength measured as intra-replicate consistency as well as increasing convergence of the phenotype (Fig. 2A). In total, 131 of 446 stimuli produced highly consistent, dose-dependent phenoprints in at least one cell type (Fig. 2B), with representation from cytokines, growth factors, chemokines, antibodies, microbial toxins, and others (Fig. S2).

**Fig. 2:**
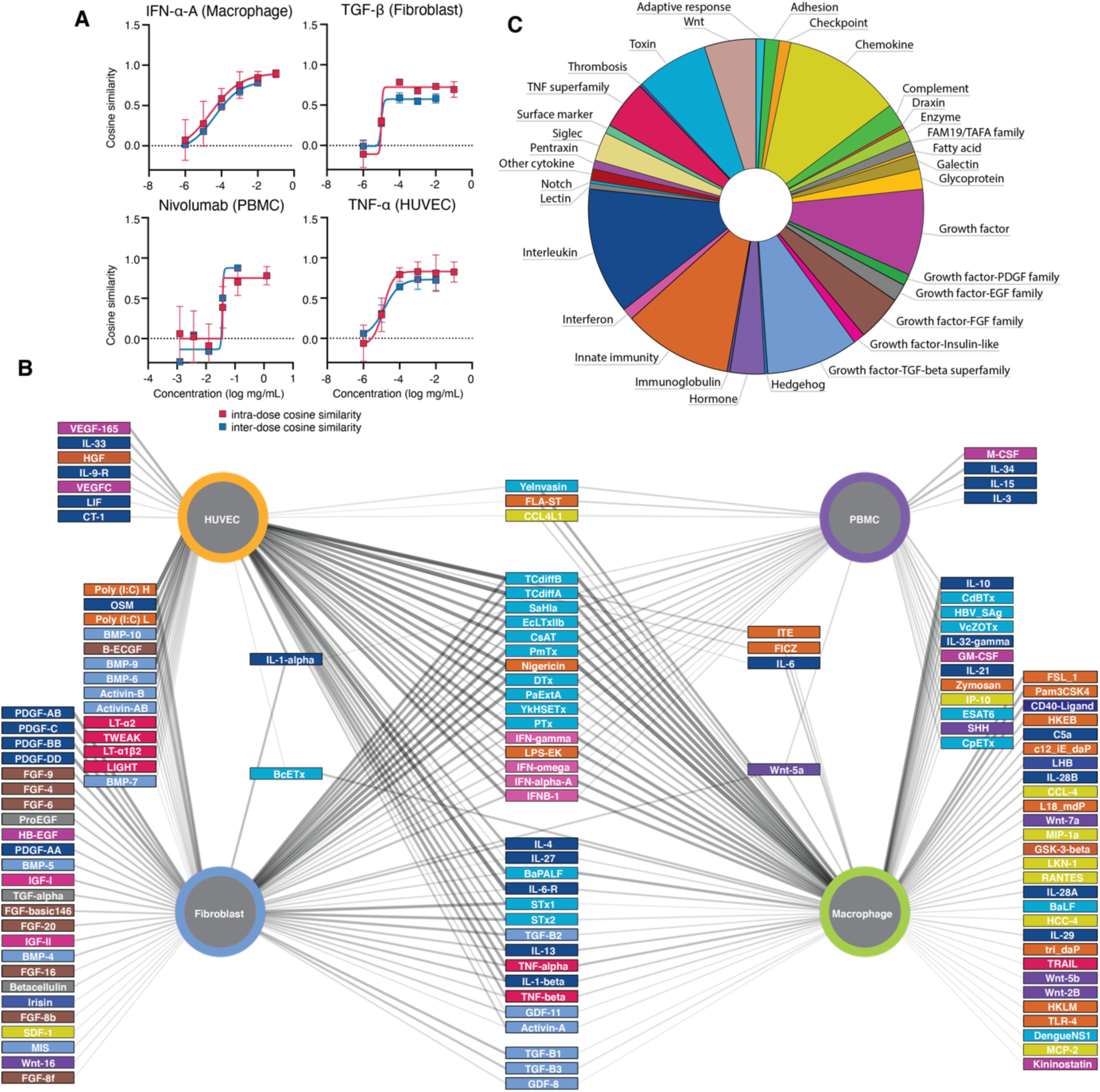
Dose dependent mapping of immune stimuli effects across multiple human cell types using phenomics. **A**. Example phenoprints for indicated immune stimuli and cells. The intra-dose cosine similarity among well replicates (n=6) for each perturbation dose, demonstrating perturbation consistency(red), and average pairwise cosine similarity between each perturbation dose (blue) and the highest dose, demonstrating phenotypic convergence. **B**. Statistically significant phenoprints induced by immune stimuli (rectangles, colored by class as in Fig. 2C) are connected by edges to the cell type (circles) in which the phenoprint was observed. Thicker edges reflect stronger interactions. **C**. Classes of all immune stimuli in immune perturbant library.

A subset of factors, such as innate stimulants like LPS, interferons, and microbial toxins (e.g. enterotoxins) produced strong dose-dependent phenoprints in all cell types tested. By contrast, other stimuli produced phenoprints only in the expected cell type: Vascular endothelial growth factor (VEGF) in endothelial cells, fibroblast growth factor (FGF) family in fibroblasts, innate stimulants and interferons in macrophages and PBMCs, and TGF-β family in both macrophages and fibroblasts^28,29^.

### Mapping high-dimensional functional relationships

To test whether phenoprints could capture known functional relationships, we applied hierarchical clustering and confirmed appropriate grouping for factors with similar function and/or structural homology, often in a cell-type-specific manner (Fig. 3A, Table S2). For example, related factors IL-4 and IL-13, and all type-I IFNs clustered in unique groups in any cell type in which a phenoprint was observed. Chemokines (such as CCL-4 and MCP-2) and innate stimulants (such as LPS and flagellin) are also readily grouped with other stimulants, but only in their appropriate cell types: macrophages and PBMCs (Fig. 3A, green/purple edges). Growth factors cluster in fibroblasts, consistent with their sensitivity to environmental cues for matrix remodeling and organ function, and ability to differentiate into other states such as myofibroblasts under inflammatory pressure (Fig. 3A, blue edges)^30,31^.

**Fig. 3:**
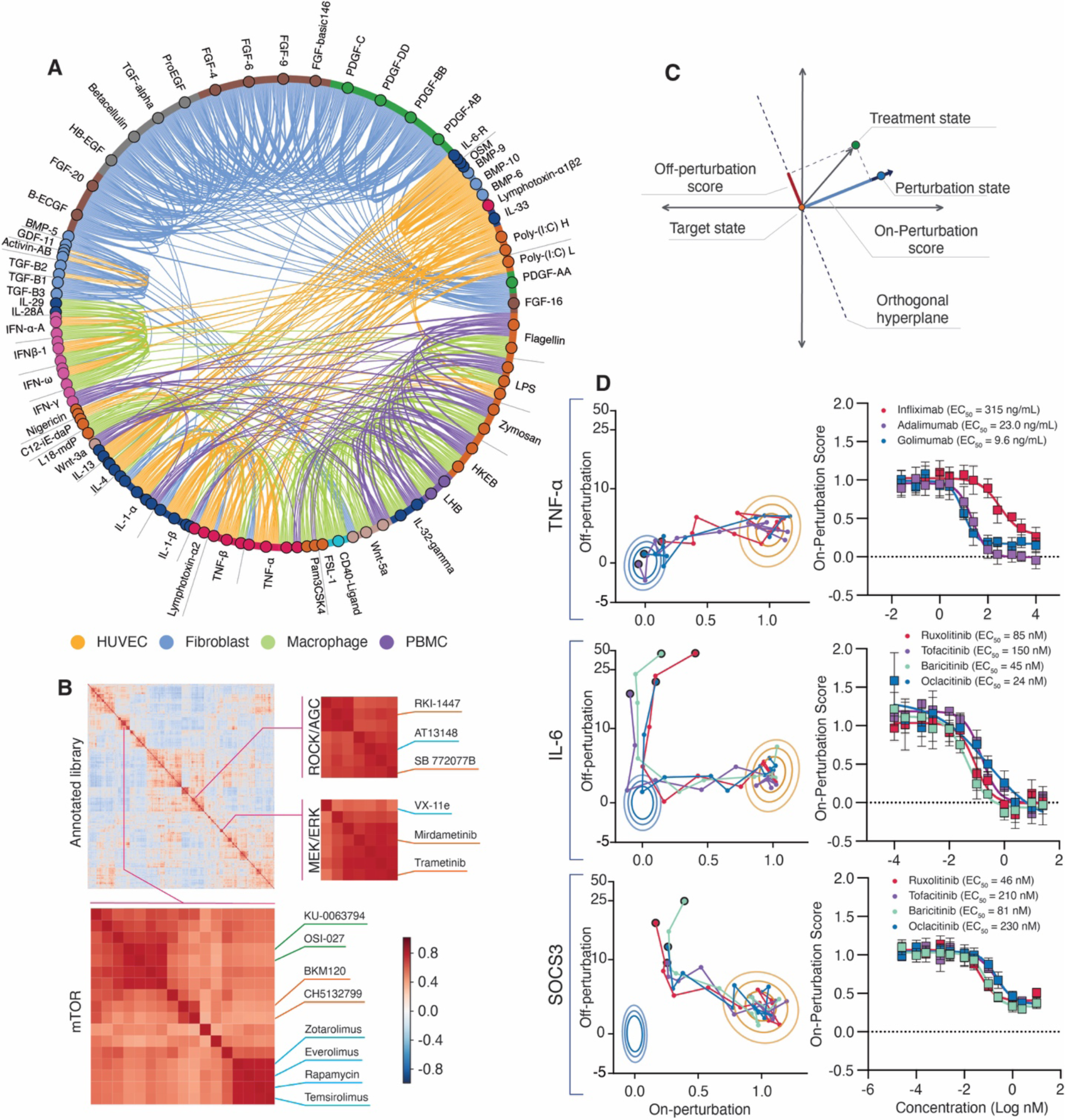
Recapitulation of known biological relationships using phenomics. **A**. High-dimensional morphological relationships in four cell contexts. Immune stimuli are arranged based on hierarchical clustering of all factors in all cell types; only factors with a statistically strong phenoprint in at least one cell type are shown. Lines between nodes are filtered for similarity and colored based on the cell type in which the association was observed. **B**. Hierarchical clustering of phenoprints resulting from HUVEC treated with an annotated compound library. Highlighted clusters pertain to small molecule inhibitors of AGC family kinases: AKT (blue), ROCK (orange), MEK (orange)/ERK (blue), and mTOR inhibitors: rapalogues (blue), PI3K inhibitors (orange) and mTORC1/2 (green). **C**. Process for reduction of high-dimensional data into two dimensions. **D**. Projections of compound response in the context of on- and off-perturbation vectors and logistical regression for on-perturbation values by drug concentration for TNF-α, IL-6 (receptor chimera) ligands, and *SOCS3* knockout in HUVEC (mean, n=6). Contour lines depict perturbation state (orange) and target state (blue) replicates at the 50, 90, and 99th percentiles, respectively.

Phenoprints also captured associations between initial stimuli and overlapping secondary effects, as in the case of clusters for activators of NFκB (TNF-α, IL1β), interferons induced by IRF3, and TLR3 ligands (Poly (I:C) variants) in endothelial cells (Fig. S1C). Double-stranded RNA motifs such as Poly (I:C) are recognized by TLR3, which signals through both NFκB and IRF3 pathways^32^.

Fine-grained distinctions were visible in clustered phenoprints. For example, the TGF-β superfamily (TGF-β proteins, growth differentiation factors, activins, and Müllerian inhibiting substance) formed a distinct cluster from other growth factor families, which included the epidermal growth factor (EGF) family (EGFs, TGF-α, betacellulin, heparin-binding EGF), platelet-derived growth factor (PDGF) family, FGF family and the insulin-like growth factor (IGF) family (Fig. S1D). Within these larger groupings, nearly structurally-identical TGF-β isoforms TGF-β1, TGF-β2, and TGF-β3 clustered more tightly than do other members of the TGF-β superfamily^33^.

Pathogen-derived toxins also revealed expected relationships. *C. difficile* toxins A and B produced cell-type-specific similarities with members of the human immune stimulant panel (Fig. S1E). These toxin phenoprints were similar to those of interferon proteins in macrophages, as well as to IL-6 superfamily members and NFκB-mediated stimulants in HUVEC. This aligns with the known dual pathology of *C. diff*. infection, involving both inflammation through macrophage activation^34^ and direct gut permeabilization effects^35^.

### Evaluating immune-relevant therapeutic candidates

Confidence in phenotypes resulting not only from the immune perturbation but those resulting from compounds in a screening library is critical for evaluation of high-throughput screens. To this end, we generated individual compound phenoprints from an annotated bioactive library, which indeed clustered by mechanism of action (Fig. 3B). For example, inhibitors of AGC kinases Akt and ROCK clustered within a super-group as did inhibitors of MEK and ERK, which is expected based on protein homology and sequential signaling, respectively. More granular sub-clusters were also observed; for example, a diverse group of inhibitors of mTOR can be sub-classified into rapalogues, PI3K inhibitors and mTORC1/2 inhibitors.

We next tested whether phenomics could uncover compounds that rescue the complex high-dimensional effect of various immune perturbants. We applied several disease-related phenoprints identified above and defined a corresponding perturbation vector in high-dimensional embedding space. Compounds that rescued on-perturbation morphology with minimal deviation in off-perturbation morphology were selected as screen hits, as they offer a potential combination of efficacy and specificity (Fig. S3A, B).

We first validated the strategy using approved anti-TNF antibodies in the context of a TNF-α phenoprint (Fig. 3D). All tested antibodies rescued the phenoprint induced by TNF-α (perturbed state) back towards the unperturbed phenoprint (target state), but infliximab was less potent relative to adalimumab and golimumab. It is known that infliximab, adalimumab, and golimumab all have similar affinities for TNF-α^36^, but only adalimumab and golimumab are effective in the absence of concomitant immunosuppression^37^. Although many factors can affect antibody performance, this finding suggests phenomics can differentiate subtleties of antibody efficacy beyond affinity for the ligand.

Next, we selected a set of clinical-stage and approved JAK inhibitors to test the ability of phenomics to model compound rescue of IL-6 signaling when cells are activated by the ligand or when disrupted by genetic modification. Tofacitinib, baricitinib, ruxolitinib, and oclacitinib reverse the phenoprint of the IL-6 receptor chimera in HUVEC in a dose-dependent manner. In addition to identifying compounds that block receptor-mediated signaling, we demonstrated compound-induced rescue of a phenotype resulting from knockout of an intracellular mediator.

Here, knockout of *SOCS3* using CRISPR/Cas9 leads to hyperactivation of JAK signalling^38^, resulting in a phenoprint that is rescued by the same set of compounds (Fig. 3D; comparison to inactives: Fig. S3C).

### Discovery of a novel small molecule inhibitor of the TGF-β phenoprint and rapid triage using phenomics

Well-designed phenotypic screening efforts benefit from being a more proximal model of the disease, and since they are not limited to pre-defined targets, offer the ability to uncover novel therapeutic pathways^39^. However, this carries the risk of investing time and resources into compounds that are later discovered to interact with known or disadvantageous pathways. To address this challenge, we leveraged the relatability of our NCE screening and annotated compound datasets to identify novel therapeutic opportunities while rapidly deprioritizing high-risk mechanistic space, as demonstrated in a screen against the TGF-β-induced phenoprint.

TGF-β, signaling through the receptor ALK5, is recognized as a primary driver of fibrosis in debilitating diseases such as idiopathic pulmonary fibrosis and renal fibrosis, as well as a significant contributor to immune exclusion in the tumor microenvironment^29^. The diversity and consistency of phenoprints recapitulating known biology suggested phenomics might discover new chemical entities (NCE) and predict mechanism of action. We therefore screened 90,000 diverse chemical starting points (Fig. S4A) against the TGF-β phenoprint (Fig. 4A). A novel compound of interest (REC-0104937) completely reversed the phenoprint at low micromolar concentrations. Further, this same compound rescued an orthogonal functional validation assay, mitigating TGF-β-induced collagen deposition with an EC_50_ of 0.763 μM (Fig. 4B). We then compared the REC-0104937 phenoprint to reference phenoprints derived from a set of 6,000 diverse, well-annotated small molecules; several known ALK5 inhibitors were highly similar (Fig. 4C). We experimentally validated the accuracy of these predictions in gold standard assays of ALK5 activity: REC-0104937 inhibited cellular p-Smad activity and cell-free biochemical ALK5 activity at 0.585 μM and 0.725 μM respectively (Fig. S4D, Fig. 4D)^40,41^. Because the advancement of TGF-β receptor inhibitors has been hampered by cardiac toxicity^42^, and because our research goals are to identify compounds acting against novel pathways, we rapidly deprioritized this compound in favor of others based on the primary screening data.

**Fig. 4:**
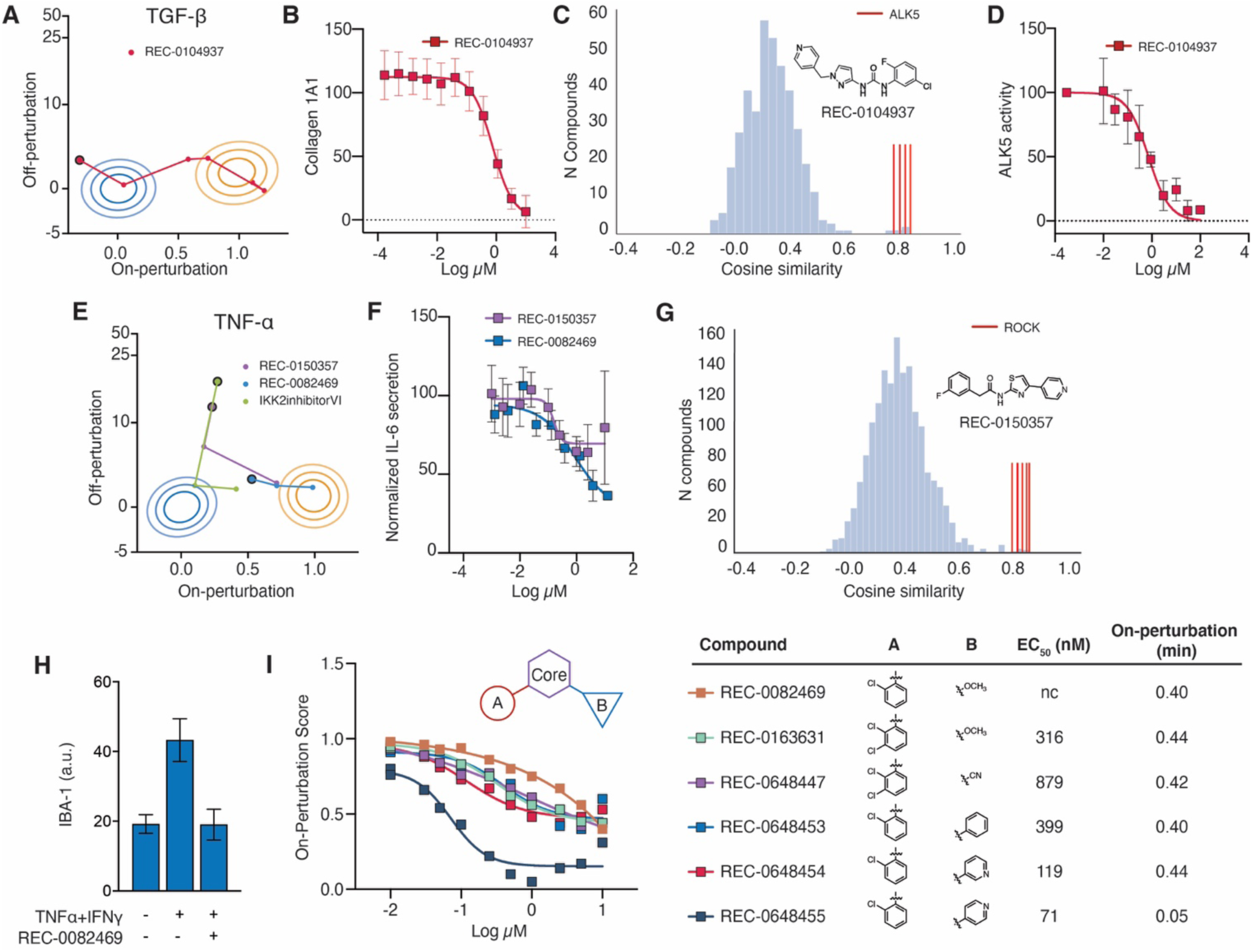
Rapid advancement of active screening compounds. **A.** Projections of compound response onto the perturbation vector for TGF-β in fibroblasts. **B**. Impact of REC-0104937 on Collagen 1A1 (n=32) expression in fibroblasts as quantified by immunofluorescence and **C**. High dimensional cosine similarity analysis of REC-0104937 compared to annotated compounds. Red bars are annotated ALK5 inhibitors. **D**. ALK5 activity in a cell-free biochemically assay (n≥2). **E**. Projections of compound response in the context of perturbation vector for TNF-α in HUVEC. **F**. IL-6 secretion (HTRF) from HUVEC treated with 1 ng/mL TNF-α in the presence of REC-0150357 and REC-0082469. **G**. Distribution of cosine similarity of phenoprints of an annotated compound library to that of REC-0150357. Red lines highlight ROCK inhibitors. Measurement of IL-6 secretion in in response to REC-0082469 and 1 ng/mL TNF-α. **H**. Rescue of TNF-α+IFN-γ-induced IBA-1 expression in microglia in the presence of REC-0082469 (n=5, mean+standard deviation). **I**. Projection of on-perturbation scores and EC_50_ values for each peripheral modification to the scaffold core (mean, n=6).

### Discovery, mechanism prediction, validation, and lead optimization of small molecules that rescue the TNF-α phenoprint in an NFκB-independent manner

Overactive TNF-α signaling is a major driver of inflammation in inflammatory and autoimmune diseases including Alzheimer’s disease^43^, multiple sclerosis^44^, and traumatic brain injury^45^, and significant benefit has been achieved with monoclonal antibody intervention^46^. We therefore sought to rescue the TNF-α phenoprint with an intervention that not only targets a novel mechanism of action, but also benefits from advantages of small molecules over existing antibodies, such as oral availability and increased central nervous system penetration.

We found 2,073 compounds that statistically altered the TNF-α phenoprint at a single dose in a 90,000-compound primary screen. We tested these for dose response and selected a subset for validation. Although suppression of TNF-α signaling through NFκB blockade is a plausible anti-inflammatory strategy, reduction of TNF-α signaling via global inhibition of NFκB leads to a challenging safety profile^47^. Two molecules (REC-0150357 and REC-0082469) rescued the TNF-α-induced phenoprint (Fig. 4E) and prevented secretion of IL-6, a marker of TNF-α stimulation (Fig. 4F) while preserving NFκB activation (Fig. S4E). By comparing the phenoprint of REC-0150357 to phenomics data from annotated compounds in prior experiments, we found strong similarity between the phenoprints of REC-0150357 and Rho kinase inhibitors (Fig. 4G). This finding was corroborated by kinase profiling, revealing ROCK1 and ROCK2 inhibition at 0.097 and 0.066 μM, respectively (Fig. S4F). During intellectual property exploration, the target was further confirmed; the scaffold was previously evaluated as a ROCK inhibitor^48^. Kinases that were inhibited to a lesser degree were CDK7 and DYRK1B (Fig. S4F). The effect of ROCK2 inhibition on TNF-α-induced inflammation is documented^51^ and similar compounds are in active development for autoimmune disease^49,50^. Therefore, we deprioritized the compound using high-dimensional primary screening data in favor of another compound, REC-0082469. The alternative scaffold also reduced TNF-α-induced IL-6 release (Fig. 4F) but did not phenotypically cluster with any of the mechanistic classes annotated in our libraries, thus enabling informed prioritization of the compound for further study. In a >400 kinase biochemical screen, the compound showed no significant inhibition of any kinase at 1 μM. To investigate the potential benefit of REC-0082469 in neuroinflammation, we explored the role of the molecule to suppress microglial activation *in vitro*. Using an immunofluorescence stain for the microglial activation marker IBA-1^51^ we confirmed that REC-0082469 reduced the activation of BV-2 mouse microglia (Fig. 4H; example images Fig. S4G). Given the potential of REC-0082469 to operate via a novel mechanism of action (MOA) we initiated a hit optimization effort initially focused on enhancing the series potency (REC-0082469 EC_50_ > 1 uM). We used phenomics as to assess our efforts to optimize potency while maintaining or improving efficacy against an unknown target; we succeeded in improving potency by more than ten-fold (REC-0648455 EC_50_ = 71 nM) and improving the on-perturbation score to 0.05 (Fig. 4I).

### Discovery of repurposing opportunities for COVID-19

The ongoing COVID-19 pandemic presents an urgent need for quick and adaptable drug discovery in the context of a complex and poorly understood disease. We leveraged our phenomics platform to screen for approved and reference (e.g., development stage antiviral) compounds that could address two key components of COVID-19 disease progression: direct effects of viral infection and the damaging effects of an unresolved inflammatory response, or cytokine storm. 1,670 and 2,913 compounds were applied to cells in the infection and cytokine storm modes, respectively, and compound rescue was evaluated with the same processing pipeline described above.

Many features of terminal COVID-19 are the result of inflammatory pressure on endothelial cells, manifesting as barrier disruption, lymphocyte recruitment, induction of blood coagulation, and acute respiratory distress syndrome (ARDS)^52^. We modeled the cytokine storm associated with late-stage COVID-19 in endothelial cells by applying cocktails of circulating proteins that mirror those from severe COVID-19^53^ patients (perturbed state) as well as healthy control patients (target state) (Table S3, Fig. S5A). We hypothesized that rescue of the perturbed state toward the target state would reveal anti-inflammatory compounds specifically relevant to the COVID-19-associated cytokine storm. Following identification of hit compounds, electric cell-substrate impedance sensing (ECIS) was employed to confirm the activity of the aforementioned compounds in an orthogonal functional model of vascular integrity challenged with the same cytokine cocktails.

Presently, JAK inhibitors have shown benefit in one non-randomized trial^54^ and represent one of the most common mechanisms being evaluated among hundreds of clinical trials active for COVID-19. In our screen against the cytokine storm phenotype, JAK inhibitors were capable of potent rescue of the severe cytokine storm phenoprint, confirming strong potential for this mechanism’s efficacy in the context of a complex immune cascade (Fig. 5A). We also identified rescue by compounds in three classes of inhibitors outside of the JAK/STAT pathway that have been less deeply explored in the context of COVID-19, including inhibitors of Syk, PI3K and c-Met. Compounds were then applied to cells with the same inflammatory cocktail and evaluated with ECIS (Fig. S5A) to inform on benefit to vascular integrity. Rescue of this orthogonal functional assay was observed for each mechanism identified in the high-dimensional assay (Fig. 4G, Fig. S5B-C).

**Fig. 5:**
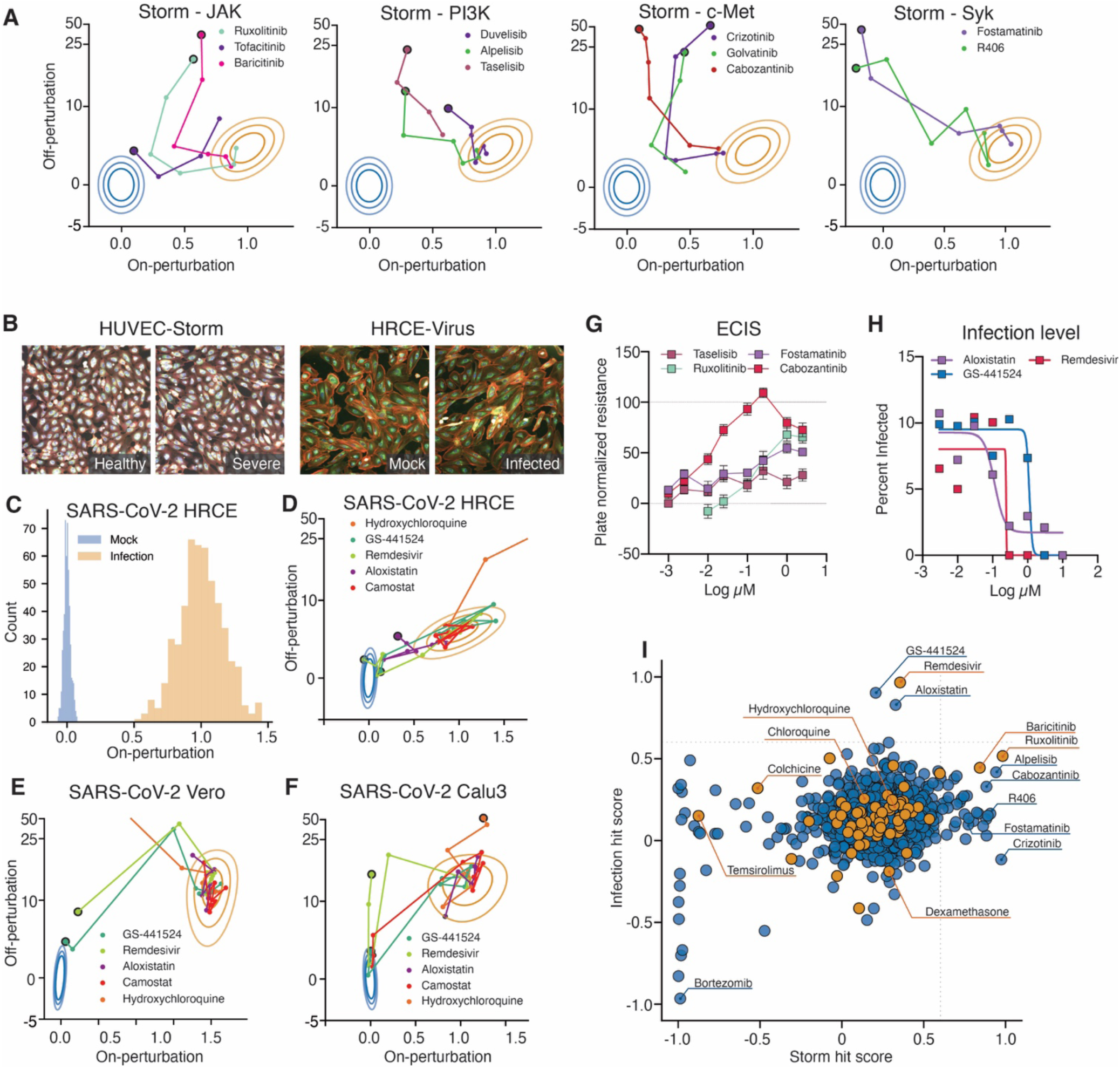
Repurposed library screening for COVID-19 using phenomics. **A**. Syk, c-Met and PI3K inhibitors rescue the severe COVID-19 specific cytokine storm high-dimensional phenoprint (perturbed state) to the healthy phenoprint (target state). **B**. Example images of target and perturbed cell populations for the cytokine storm and SARS-CoV2 viral models. **C**. Infection of HRCE yielded a phenoprint against the mock-infected target population with an assay z-factor of 0.43 for the separation in on-perturbation score for the mock and infected populations. **D-F**. Projections of compound response in the context of the perturbation vector generated in SARS-CoV-2-infected HRCE, Vero, and Calu3 cells. Off-perturbation values clipped at 50 for visualization. **G**. Compound impact on endothelial barrier function as quantified by ECIS assay. Values are normalized from 0 (cytokine storm cocktail-treated wells) to 100 (mock-treated wells). Data was averaged over a 12-minute window at hour 12 of ECIS measurement to visualize concentration response curves for the indicated compounds. **H**. Infection rate as determined by SARS-CoV-2 nucleocapsid antibody staining of infected HRCEs treated with the denoted compounds. **I**. Plot of efficacious molecules by hit-scores in SARS-CoV-2 HRCE assay vs cytokine storm assay. Orange circles denote molecules registered in interventional COVID-19 clinical trials at the time of submission. Dotted lines presented as a visual guide depicting a hit score of 0.6.

We next developed a model of SARS-CoV-2 infection to screen for repurposable compounds acting directly against viral targets or on host pathways. To define the model, we evaluated the effect of SARS-CoV-2 infection in multiple cell types, of which three resulted in robust phenoprints as compared to either mock infected or inactivated virus control populations: Calu3 (a lung adenocarcinoma line), Vero (an immortalized interferon-deficient African green monkey kidney line^55^), and primary Human Renal Cortical Epithelium (HRCE) (Fig. 5C, Fig. S6D). We confirmed active infection with SARS-CoV-2 nucleocapsid antibody staining and quantification of productive viral replication (Fig. S6A-C). We reasoned that a primary human cell type would be most directly translatable^56^ to human pathology, especially from tissues demonstrated to be directly infected by SARS-CoV-2^57^, and thus conducted a screen of 1660 compounds against the HRCE phenotype, while testing a limited subset of those compounds in Vero and Calu3 cells.

The majority of compounds currently under evaluation^58^ in human clinical trials for COVID-19 showed no or weak efficacy in the HRCE model^56^. However, in these screens remdesivir and its metabolite, GS-441524, demonstrated strong efficacy and aligned with potency described in the literature (EC_50_ of 100 nM and 2 μM, respectively) (Fig. 5D)59,60. Remdesivir is a nucleoside analog that directly interferes with the viral-RNA-dependent RNA polymerase to inhibit viral replication and, importantly, successfully reduced recovery time for treated patients in clinical trials^61^ announced after our data and analysis was publicly released. Further illustrating the predictive capacity of the model, two other antivirals, lopinavir and ritonavir were not found to be efficacious and were later discontinued in clinical testing for COVID-19^62^.

Additionally, aloxistatin (E64d), an irreversible cysteine protease inhibitor initially developed for muscular dystrophy^63^, also demonstrated suppression of the viral phenoprint in HRCEs (EC_50_ of 40 nM). Recent studies have confirmed that cathepsin L, a cysteine protease, is required for SARS-CoV-2 entry in some cell types, and aloxistatin treatment significantly reduced entry of SARS-CoV-2 pseudovirions^64,65^. We then tested a subset of these antiviral compounds in additional cell types, Vero and Calu3, and found aloxistatin did not rescue in these models. However, another protease inhibitor, camostat mesilate, was efficacious in the Calu3 model (EC_50_ of 260 nM), but not the Vero or HRCE models (Fig. 5D-F, Fig. S6E). Camostat inhibits TMPRSS2, which was recently shown to be required for SARS-CoV-2 entry in human airway cells^66^. Similar to findings in recent clinical trials^67,68^, we found chloroquine and hydroxychloroquine to have no benefit in the HRCE or Calu3 models; however, they showed modest benefit in Vero cells (EC_50_ of 730 nM and 1.65 μM, respectively) with very high off-perturbation activity (Fig. 5D-F). Overall, compound efficacy in human cell types was poorly recapitulated in Vero cells (Fig. 5D-F, Fig. S6F). Taken together these findings suggest that SARS-CoV-2 entry protease inhibitor activity varies across cell type and species; however, remdesivir and GS-441524 show strong rescue of the viral phenoprint in all cell types tested.

We identified JAK inhibitors ruxolitinib and baricitinib as efficacious in both viral and cytokine storm models (Fig. 5A, 5i, Fig. S6G). However, we found that high concentrations of these compounds led to increased infection in HRCE cells (Fig. S6H). Suppression of interferon production is a known component of SARS-CoV-2 infection at a low multiplicity of infection^69^. It is unclear however, what effect additional interferon suppression would have *in vivo*, especially at higher viral loads, warranting investigation into alternative mechanisms of cytokine storm suppression, such as PI3K or c-Met inhibition. Notably, bortezomib exhibited poor performance in both assay modes, is reported to impair endothelial cells in inflammatory contexts^70^, and also enhances susceptibility to viral infection^71^, particularly coronaviruses^72^.

## DISCUSSION

Biology is massively complex and highly networked, but the tools to explore and discover novel biology and develop medicines have until recently relied on simple, univariate measurement. The genomic revolution yielded a taste of what is possible if high-dimensional biology can be scaled by massively increasing the rate of understanding of the role of thousands of genes in human biology and disease. Following this advancement, several new high-dimensional approaches have been developed to add clarity to complex functional relationships and discover new therapeutics, but these are hindered from high-throughput screening application by engineering and cost bottlenecks. We present here early data from our experience using phenomics (Fig. 1) as one strategy to accelerate drug discovery.

We first established that diverse immune biology and pharmacology can be detected and discriminated using phenomics (Fig. 2). These data also reveal that phenomics is not simply a classification technology: deep quantification of rich, multi-parametric signal and assessment of dose response is achievable, enabling comparisons and clusterings of diverse biological perturbants alone, or in combination across diverse cell types (Fig. 3). Diving more deeply on just two of the 131 immune phenoprints we uncovered that are suitable for drug screening, we explored 90,000 new chemical entity starting points in the context of TGF-β- and TNF-α-induced phenoprints (Fig. 4). Among hits in each context, prediction of ALK5 and ROCK inhibition allowed us to rapidly shift resources to higher priority hits. In particular, we focused our efforts on a suppressor of the TNF-α phenoprint with a high potential to potentially be active against a novel, but as of yet unknown, target. Further, we drove medicinal chemistry work against this unknown target(s) using phenomics, demonstrating a 10-fold increase in potency while also increasing the magnitude of rescue.

The application of phenomics can be extended to more complex disease-causing perturbations as well: the platform was rapidly adapted for the characterization and exploration of actionable therapies in the context of a novel and poorly understood disease, COVID-19. Within 28 days of initiating the project, we identified hits through high throughput chemical screens against COVID-19 cytokine storm and SARS-CoV-2 infection in the relevant tissue types, without the need to develop cytokine- or virus-specific reagents and assays. We demonstrated that a handful of drugs currently in clinical trials strongly modulate the infection model (e.g. remdesivir), the cytokine storm model (PI3K inhibitors) or both (JAK inhibitors), prior to their clinical trial results becoming available. Conventional antiviral research relies heavily on univariate assays that measure attributes like cell death or expression level of one protein. Using a single platform, we found not only conventional antivirals, but also compounds with unconventional effects on disease-associated host pathways such as inflammation.

In the SARS-CoV-2 model remdesivir and its analog, GS-441524 demonstrated efficacy in all cell models tested. Unfortunately, remdesivir is dosed via an intravenous route, typically in an inpatient setting and a time at which cytokine storm may be primarily responsible for the pathology (wherein remdesivir had no unexpected efficacy in our cytokine storm model). SARS-CoV-2 is able to use a variety of receptors to facilitate cell entry, with receptor specificity by cell type apparent in our data: aloxistatin (E64d), inhibiting the cathespin-mediated entry pathway, and camostat, inhibiting the TMPRSS2-mediated pathway, each demonstrated strong response in HRCE and Calu3 cells respectively. Nevertheless, pseudovirus entry assays^66^ have shown that even in cells with both pathways active, modulating a single pathway still quantitatively reduces viral infection load. Further study of the proportional activity of each pathway in relevant human tissues may be warranted. As aloxistatin is orally bioavailable, simply and inexpensively synthesized, and has a relatively strong safety profile based on chronic treatment of muscular dystrophy patients in a phase 3 trial, it deserves further study for COVID-19, with the expectation that early treatment in the course of infection may be most efficacious.

In our model of more advanced COVID-19 symptoms driven by cytokine storm, JAK inhibitors were noteworthy rescuers of the inflammatory phenoprint and moderate rescuers of the viral phenoprint at low concentrations. Due to oral bioavailability and the safety profile of acute treatment they are excellent candidates for repurposing. However, we observed that JAK inhibitors enhanced cellular infection of SARS-CoV-2 at higher concentrations, suggesting an effect on interferon signaling, a possible clinical liability that should be closely monitored during trials. Supporting this finding and underlining the importance of identifying diverse options to address cytokine storm, JAK inhibitors are known to increase the prevalence or severity of other viral infections including herpes zoster, JC virus, and hepatitis B^73–75^. This study also identified alternative mechanisms of action which have been much less deeply considered in the context of COVID-19, such as certain Syk inhibitors, c-Met inhibitors and PI3K inhibitors. Such molecules could be critical additions to remdesivir therapy in severe patients. This work and the recent success of dexamethasone in clinical trials for COVID-19 also identified a key limitation of our current phenomics approach: when studying a cell type in isolation, phenomics surfaces compounds that act via cell-autonomous mechanisms^76^. Compounds that intervene in multicellular processes might be revealed by development of co-culture models.

Taken together, our results demonstrate that systems-level modeling and drug discovery is achievable using a single phenomics platform. First, this approach simplifies and extends the ability to work across many disease models rapidly because assay development work for any new model is minimized. Second, this work partially overcomes a historical limitation of phenotypic screening, predicting mechanism of action, by relating the high-dimensional phenoprint of hit compounds to those of reference molecules. Finally, we show the potential of this platform in optimizing NCE compounds through medicinal chemistry in a high-dimensional, target-agnostic manner. Unlike other high-dimensional approaches, the relatively inexpensive nature of these image-based assays allows them to be scaled to levels of throughput comparable to more traditional low-dimensional screening modalities. In the hopes that it will be valuable to others, we have made images and embeddings from HUVEC treated with the immune perturbant library, and from our COVID-19 primary screens (both infection and cytokine storm) available online (including raw image data, metadata, and deep learning embeddings from images) at rxrx.ai/rxrx2 and rxrx.ai/rxrx19, respectively.

Beyond the applications described here, the modular nature of this phenomics platform enables rapid adaptation to different libraries of immune stimulants, antibodies, or other large molecules, and incorporation of additional cellular contexts like co-culture models. In future work, comparisons of hits to phenoprints associated with knockout of each gene in the genome (achieved by arrayed whole-genome CRISPR knockout) may further expand our ability to predict mechanisms beyond those represented within our annotated small molecule library, bridging a key gap in phenotypic screening. Critically, these data can be related over time and across disparate research programs—supporting the creation of large biological image datasets for deep-learning applications^77,78^ that will accelerate drug discovery and yield functional maps of human cellular biology.

## MATERIALS/METHODS

### Cells

**HUVEC:** Human umbilical vein endothelial cells (Lonza, C2519A) were cultured according to manufacturer’s recommendations in EGM2 (Lonza, CC-3162). **NHLF:** Normal Human lung fibroblasts (Lonza, CC-2512) were cultured according to manufacturer’s recommendations in FGM2 (Lonza: CC-3131, 4126) and used at passage four for all assays. **PBMC:** Peripheral blood mononuclear cells (PBMCs), from healthy donors, were prepared from fresh (< 24 hours old) leukopaks (STEMCELL Technologies Inc., Catalog # 70500). Following 3 120 rcf washes (brake off) for platelet removal, the samples were processed by EasySepTM RBC Depletion Reagent in accordance to manufacturer’s instructions (STEMCELL Technologies Inc., Catalog #18170). Following isolation, PBMCs were pelleted (300 rcf) and resuspended in cryo-preservation medium (CryoStor® CS10 Freeze Media, BioLifeSolutions Inc., Part #: 210102) for long term storage. **Macrophage**: Macrophages were derived from either cryo-preserved or freshly isolated PBMCs. Monocytes were enriched by plastic adherence, first seeding PBMCs in serum free RPMI-1640 medium followed by 1.5 hours of incubation and washed 2x with PBS. Cells were incubated for 3 d in complete medium (RPMI+ 10% heat inactivated FBS, 25 ng/mL M-CSF, 10 ng/mL IL-10). Media was replenished after 3 d with one third of the conditioned medium and 2/3 fresh complete medium. Monocyte Derived macrophages (MDMs) were harvested from each vessel after an additional 3 d using ACCUTASE following manufacturer’s instructions (Thermo - A1110501). MDMs were pelleted (300 rcf) and resuspended in cryo-preservation medium. **HRCE**: primary human renal cortical epithelial cells Lonza (CC-2554) were propagated at 37°C with 5% CO2 in EpiCM, (ScienCell # 4101) supplemented with Epithelial Cell Growth Supplement (EpiCGS, ScienCell #4152). **Vero**: an immortalized african green monkey kidney (ATCC CCL-81) were propagated at 37°C with 5% CO2 in Eagle’s Minimum Essential Medium (EMEM) supplemented with 10% FBS. **Calu3** human lung adenocarcinoma line (ATCC HTB-55) were propagated at 37°C with 5% CO2 in EMEM supplemented with with 10% FBS. **BV-2**: murine microglial cells (ICLC ATL03001, Ospedale Policlinico San Martino) were propagated at 37°C with 5% CO2 in RPMI media + 10% Heat Inactivated FBS.

### Preparation of stimulant library

Immune stimuli (Table S1) were solubilized in sterile phosphate buffered saline (PBS) containing 0.1% BSA (Sigma cat.# A1595-50ML) to make stock solutions of .04 mg/mL in Echo-qualified 384W low-dead volume source plates. Source plates were stored at −80°C until use.

### Generating high-dimensional phenotypes

Cells were seeded into 1536-well microplates (Greiner, 789866) via Multidrop (Thermo Fisher) and incubated at 37C in 5% CO_2_ for the duration of the experiment. Immune stimuli or virus were added 24 hours post-seeding (HUVEC, Macrophage, Fibroblast) or 1 h (PBMC). Treatments were randomized across treatment plates with a 6-log range of immune stimuli (typically 0.001-100 ng/mL) at 6 replicates each with acoustic transfer (Echo 555, Labcyte) and incubated 37°C for 24 or (complete immune stimuli panel) or 48 h (for PBMC with pembrolizumab or nivolumab). Active SARS-CoV-2 was added via multidrop 24 hours post seeding of the specified cell type. Plates were stained using a modified cell painting protocol^29^. Cells were treated with mitotracker deep red (Thermo, M22426) for 35m, fixed in 3-5% paraformaldehyde, permeabilized with 0.25% Triton X100, and stained with Hoechst 33342 (Thermo), Alexa Fluor 568 Phalloidin (Thermo), Alexa Fluor 555 Wheat germ agglutinin (Thermo), Alexa Fluor 488 Concanavalin A (Thermo), and SYTO 14 (Thermo) for 35 minutes at room temperature and then washed and stored in HBSS+0.02% sodium azide. Live-virus experiments omitted the mitochondrial stain due to operational constraints of the biosafety level environment.

### Compound screen and imaging

One hour prior to addition of the immune stimulant or 18 hours prior to the addition of virus, cells were treated with compound via acoustic transfer (Echo 555, Labcyte). Primary screening of New Chemical Entity libraries was performed at 10 or 30 μM with concentration-response confirmation spanning 100 nM to 30 μM in half-log steps. SARS-CoV-2 screening was completed in dose response in half-log steps between 10 nM to 3μM. After 24 h incubation (or 96 hours post viral infection), plates were imaged using Image Express Micro Confocal High-Content Imaging System (Molecular Devices) microscopes in widefield mode with 20X objectives. Four sites per well were acquired with 6 channels per site. The following bandpass filters were used to visualize the channels: FF409/493/573/652, FF459/526/596, FF01-432/515/595/730-25, FF01-475/543/702, and FF01-600/37/25.

### Kinase analysis

Kinase profiling was performed using a KINOMEscan^™^ panel of 97 or >400 kinases at Eurofins-DiscoverX (San Diego, CA). Targets exhibiting > 50% inhibition were followed by KdElect^®^ analyses for DYRK1B, CDK7, ROCK2 and ROCK1 to determine IC_50_’s. Data presented as mean and standard deviation.

### CRISPR gene knockout

Alt-R Crispr-Cas9 reagents were purchased from Integrated DNA Technologies, Inc. (IDT) and prepared following the manufacturer’s guidelines and protocols (Alt-R CRISPR-Cas9 crRNA, Alt-R CRISPR-Cas9 tracrRNA cat #1072534, Alt-R S.p. Cas9 Nuclease V3, cat #1081059, and Alt-R Cas9 Electroporation Enhancer, cat #1075916). Alt-R CRISPR-Cas9 crRNA was duplexed to Alt-R CRISPR-Cas9 tracrRNA and then combined with Alt-R S.p. Cas9 Nuclease V3, following IDT guidelines, to form a functional CRISPR-RNP complex. This CRISPR-RNP complex was transfected into cells using the Lonza 4D Nucleofection system and standard protocols with proprietary modifications, or with a proprietary lipofection-based process for high-throughput application. Alt-R Cas9 Electroporation Enhancer was included into the nucleofection reactions to enhance transfection efficiency following standard guidelines from IDT.

### Image featurization for phenomic analysis

All images were uploaded to cloud storage and featurized by embedding them with a trained neural network using Google Cloud Platform. This network is based on the convolutional neural network DenseNet-161^30^. We adapt this network in the following ways. First, we change the first convolutional layer to accept image input of size 512 × 512 ⨯ 6. Like DenseNet-161, we use Global Average Pooling to contract the final feature maps, which in our case are tensors of dimensions 16 ⨯ 16 ⨯ 2,208, to a vector of length 2,208. However, instead of following immediately with a classification layer, we add two fully-connected layers of dimension 1,024 and 128, respectively, and use the 128-dimensional layer as the embedding of the image. The weights of this network were learned by adding two separate classification layers to the embedding layer, one using softmax activation and the other using ArcFace^78^ activation, which were simultaneously optimized by training the network to recognize perturbations in the public dataset RxRx1^79^ and in a proprietary dataset of immune stimuli in various cell types. Due to operational constraints of the BSL-3 assay conditions, a modified assay protocol lacking one image channel was used for the live-virus experiments. To accommodate this change, we trained a separate network of the same basic architecture that used only five input channels and one fully-connected final layer of dimension 1,024.

### Phenomic Analysis

Immune stimuli phenoprints were observed by calculating the mean embedding of all but one biological replicate, finding the angle between that average and the held-out replicate well, and repeating this process for every replicate to find the average cross-validated angle for that perturbation. Statistical significance of these phenotypes was determined by comparing their similarity at high dose against a distribution of similarities between embeddings of images of untreated cells. We used the Benjamini-Hochberg multiple tests correction with a 5% false discovery rate and considered phenotypes acceptable if they had a corrected p-value<0.05 in two independent experimental batches.

The similarity between a pair of immune stimuli was determined by calculating the cosine similarity between all pairs of embeddings of one immune stimulant at high concentration with the embeddings of another immune stimulant at high concentration, and testing whether the mean of this pairwise-similarity distribution was significantly different from zero using a one-sample t-test and employing the Benjamini-Hochberg multiple tests correction with a 5% false discovery rate. Only significant pairs are used in this paper, and the means of their pairwise-similarity distributions are the values reported in the figures.

For small molecule screens, post-processing of the embedded images included normalization to remove inter-plate variance, PCA to reduce the feature space, and anomaly detection to remove outliers from the control populations. The vector pointing between the barycenters of the untreated and perturbed conditions was computed, and the embedded image vectors were decomposed into the signed scalar projection (the on-perturbation score) and the scalar rejection (the off-perturbation score) with respect to this vector. These scores were normalized so that the mean on-perturbation score was 0 for the untreated condition and 1 for the perturbed condition. Separation of the untreated and perturbed conditions along the on-perturbation axis was assessed by Z-factor.

For compound MOA inference, cosine similarities were computed between the embeddings of an NCE compound and the set of embeddings of a compound library annotated for MOA, and significantly large similarities (relative to the distribution of similarities of pairings of annotated compounds with the NCE compound) were reported.

### Cytokine storm cocktails

For the cocktail representing severely affected patients, top concentration of the most abundant protein, CXCL-10 was selected to be 200 ng/mL based on a practical screen concentration and previously identified phenotypes for this factor. All other proteins were prepared at appropriate concentrations relative to CXCL-10. Cocktails representing healthy patients and those with moderate disease severity were prepared with each concentration relative to the severe cocktail.

### Iba1 Immunofluorescence assay

BV-2 microglia were thawed from liquid nitrogen and plated at 2500 cells per well in 384-well PDL/collagen-coated plates (Greiner #781866). The next day, the cells were treated with compound first, followed an hour later by stimulant (recombinant murine TNF-α+IFN-γ (Peprotech), 200 μg/mL in 0.1% low endotoxin BSA/PBS). Twenty-four hours after treatment, BV-2 microglia were fixed and stained for the microglial activation marker Iba1. Briefly, cells were fixed with 4% PFA for 15 min at room temperature. Primary antibody solution was added to a 1:100 final dilution (Iba1 antibody Abcam cat #ab5076) incubated overnight at 4°C. After overnight incubation with primary antibody cells were washed with PBS, and secondary antibody solution was added (AlexaFluor 488 donkey anti-goat IgG, 1:1000 final dilution. Invitrogen cat# A11055). Cells were then incubated for 1 hr at room temperature, protected from light. Following secondary antibody incubation cells were washed and the plate was sealed for imaging, which was performed on an Image Express Micro Confocal High-Content Imaging System (Molecular Devices). Data and error presented as mean and standard deviation.

### HTRF

Quantitative measurement of cytokines in supernatants obtained from cultured cells was performed by using a homogenous time-resolved fluorescence assay (HTRF, Cisbio). IL-6 HTRF assays were performed in accordance with the manufacturer’s protocol. Briefly, cells were seeded and after 24 h treated with compound over 8 concentrations and 5 replicates each. After an additional 24 h, supernatant was collected from assay plate and appropriate sample dilutions and standards were made and dispensed into barcoded labeled 384-PerkinElmer ProxiPlates (cat# 6008280). After the recommended incubation time, the plate was read using an Envision^®^ 2105 microplate reader (PerkinElmer). Data and error presented as mean and standard deviation.

### NFκB translocation assay

NFκB reporter cells (TR860A-1, System Biosciences) were seeded at 4000 cells per well in 384 well imaging plates (781948, Greiner). Compounds were added at at least 4 replicates per concentration followed by 1 ng/mL TNF-α after 1 h. Plates were imaged (GFP channel) once every 3 h via incucyte (Sartorius). Data for Integrated intensity over at the 16 h time point is presented as mean and standard deviation, significance analyzed with 2-way ANOVA.

### pSmad assay

Normal Human Lung Fibroblasts (NHLF, Lonza) were plated in 1536-well plates (Greiner) at 0.25 × 10^6 cells/mL in FGM-2 (Lonza). After 24 hours, the media was replaced with FBM media (Lonza) and incubated for 24 hours. Cells were treated with compounds of interest in a 11-point dose response curve at 32 replicates per concentration using an acoustic liquid handler, and incubated for 1 hour at 37°C, 5% CO2. Cells were then treated with 1uL of 11ng/mL TGF-β1 (R&D Systems), for a final concentration of 1ng/mL. Cells were incubated for 30 minutes at 37°C and 5% CO2. Cells were fixed with 4% PFA, blocked for 1 hour with 1% BSA/0.1% Triton X-100/PBS, and then stained for pSMAD (Cell Signaling, 1:800). After an overnight incubation at 4°C, cells were stained with AlexaFluor 647 (Thermo, 1:1000) and Hoescht (Thermo, 1:5000) for two hours at 25°C, washed with PBS twice, and imaged on an Image Express Micro Confocal High-Content Imaging System (Molecular Devices). Images were analyzed with CellProfiler to observe nuclear translocation of pSMAD2. Data and error presented as mean and standard deviation.

### Collagen expression

Normal Human Lung Fibroblasts (NHLF, Lonza) were plated in 1536-well plates (Greiner) at 0.25 × 10^6 cells/mL in FGM-2 (Lonza). Cells were treated with compounds of interest in a 11-point dose response curve at 32 replicates per concentration using acoustic transfer (Echo 555, Labcyte), and incubated for 1 hour at 37°C, 5% CO2. Cells were then treated with 1uL of 11ng/mL TGF-β1 (R&D Systems), for a final concentration of 1ng/mL using Multidrop (Thermo Fisher). Cells were incubated for 96 hours at 37°C and 5% CO2. Cells were fixed with 4% PFA, blocked for 1 hour with 5% BSA/0.2% Triton X-100/PBS, and then stained for Collagen I (Cell Signaling, 1:500). After an overnight incubation at 4°C, cells were stained with AlexaFluor 750 (Thermo, 1:1000), CellMask Orange (Thermo, 1:5000) and Hoescht (Thermo, 1:5000) for two hours at 25°C, washed with PBS twice, and imaged on an Image Express Micro Confocal High-Content Imaging System (Molecular Devices). Images were analyzed with CellProfiler. Data and error presented as mean and standard deviation

### Electric Cell-substrate Impedance Sensing (ECIS)

Prior to use, 96-well ECIS plates (Applied Biophysics, 96W20idf PET) were pre-treated with 10mM L-cysteine (Sigma-Aldrich, C7352-25G) and then coated with fibronectin (gibco, PHE0023). Human Umbilical Venous Endothelial Cells were plated in the fibronectin coated 96-well ECIS plates at 55,000 cells/well in EBM-2 (Lonza cc-3156) +EGM-2 (Lonza, cc - 4176). Cells were allowed to settle at room-temperature for 1 hour and then incubated for 24 hours at 37^°^C, 5% CO_2_. Following incubation, the plates were placed on the ECIS readers for 1 hour to establish baseline resistance. Cells were then treated in a 9-point dose response curve at 4 replicates per concentration using acoustic transfer (Echo 555, Labcyte) and returned to the incubator for 1 hour. Following this, cells were treated with the cytokine storm cocktail (Table S3). Resistance was measured for 24 hours following this. The assay window was defined as the time range with the greatest observable difference in membrane resistance between empty and disease control (approximately 6 hours following cocktail addition). Resistance was normalized to each plate and graphed as a dose-response curve where 1 and 0 correspond to health and disease controls, respectively.

### SARS-CoV-2 Nucleocapsid staining

After staining and imaging to establish high dimensional phenotypes, plates were rinsed once with Wash Buffer (1xHBSS + 0.02% sodium azide) before incubating with primary antibody raised against SARS-CoV-2 nucleocapsid protein for 60 mins at RT (Sino Biological catno. 40588-T62, 1:1000 dilution). Media was evacuated from wells by inverted centrifugation, and secondary antibody was added and incubated another 60 minutes at RT (Thermo Scientific catno. A31573, 1:2000 dilution). Primary and secondary antibodies were diluted in Stain Base media (1xHBSS, 1% BSA, 0.3% Triton-X 100). Plates were washed one final time using inverted centrifugation and Wash Buffer before imaging as described above. Data presented as mean value.

### Calculation of SARS-CoV-2 infection Rate

Cell-level image segmentations and per-cell log-mean nucleocapsid staining intensities were calculated using standard image segmentation techniques (CellProfiler^80^). These intensities were normalized by plate with respect to log-mean-intensity in the mock control cells. To adjust for optical effects that changed the background fluorescence level in infected wells, a Gaussian mixture model was used to align the lowest peak in log-mean-intensity across well conditions. Cells were estimated to be infected if they exhibited an adjusted log-mean-intensity above the 99th percentile of intensity for the control cell population. This estimate of number of infected cells was used to compute a fraction of cells infected, which was adjusted to account for the 1% of uninfected cells expected to be above the 99% threshold.

### SARS-CoV-2 propagation and controls

The USA-WA1/2020 strain of SARS-CoV-2 was propagated in Vero cells. Cells were grown in standard tissue culture flasks (60% confluence) and were infected at a multiplicity of infection (MOI) of 0.001, in EMEM + 2% FBS and 50 g/mL gentamicin, incubated at 37°C with 5% CO2 for 5 days. Supernatants containing virus were removed from these cultures, spun down to remove cellular debris and stored at −20°C until use. Viral titers were determined through standard tissue culture infectious dose 50% (TCID50) methods, where cytopathic effect (CPE) on Vero 76 cells was measured by visual observation under a light microscope.

To create a suitable control with inactivated virus, SARS-CoV-2 was irradiated with a UV lamp for 10 or 20 minutes. Viral inactivation in this sample was verified using visual CPE on Vero cells, where undetectable level of active virus was observed. An additional “mock” control was created using conditioned media preparations generated from uninfected Vero 76 cells grown in 2% FBS in EMEM for five days. Cellular debris were removed through centrifugation and the supernatants were frozen at −80°C until use.

All experiments using SARS-CoV-2 were performed using Biosafety Level 3 (BSL-3) containment procedures at partner facilities including one at Utah State University. Data and error presented as mean and standard deviation.

### ALK5 biochemical assay

A 1536-well plate was pre-treated with compounds, at 13 concentrations (ranging from 0 to 100 ng/mL) with at least 2 replicates of each concentration, and a reaction mix containing 15 ng ALK5 (ThermoFisher), using Poly 4:1 as substrate was added to the plate. The reaction was started by the addition of 20 μM ATP, the plate was mixed on a plate shaker (500 RPM for 2 min) and the reaction allowed to incubate at room temp for 60 min. The reaction was terminated by the addition of ADP-Glo Reagent (Promega V9102) (40 min incubation) and Kinase Detection Reagent (30 min incubation). Luminescence was captured using an EnVision XCite plate reader. Data presented as mean and standard deviation.

### Small molecule structural analysis

All assessments of compound diversity and similarity were performed in DataWarrior (openmolecules.org). Neighbors were determined to be at 83% or greater similarity as determined with the SkelSpheres descriptor.

## Supporting information

Supplementary table 1_Immune perturbants

Supplementary table 2_Cellular phenotypes and classes

Supplementary table 3_Cytokine storm cocktails

## Data availability

Tables containing metrics for immune stimulant phenotypes in each cell type are provided in the Supplementary Materials. Underlying images, metadata, and deep learning embeddings for soluble factor perturbations in HUVEC and all primary screens in primary cell types for COVID-19 virus and cytokine storm screening have been made available at rxrx.ai. An interactive server containing drug response projections, hit scores, and structures for COVID-19 screening data has been made available for custom search at covid19.rxrx.ai.

## Code availability

The code underlying this report leverages proprietary algorithms for image processing, data standardization, outlier detection, and compound efficacy scoring. As such the code underlying this report will not be made available. Instead, much of the output of these algorithms is provided in the provided Supplemental Tables.

Virus graphic in Fig. 1 adapted from an image by Desiree Ho for the Innovative Genomics Institute and is licensed under a Creative Commons Attribution-NonCommercial-ShareAlike 4.0 International License

No replicates presented were repeat measurements of the same biological conditions.

## Acknowledgements

We would like to acknowledge members of the Recursion team, including Scott Ackler, Anna Adamson, Jorge Aguilera, August Allen, Lawrie Allred, Emelly Alvarado, Teresa Anderson Myers, Charles Baker, Ryan Baker, Benjamin Banowsky, Celeste Belletire, Kathleen Bennett, Daniel Berger, Ashish Bhandari, Andrew Blevins, Chad Bradford, Katie Brown, Kevin Brown, Matthew Burbidge, Emmanuel Cardenas, Nicholas Cernek, CJ Christensen, Brooke Clark, Cossette Clemens, David Compton, Hannah Cook, Jacob Cooper, Rachel Crane, Timothy Dahlem, Aaron Daniels, Madison Davis, Quincy Davis, Shawndi Ellis, Joel Ellis, Aisha Fairclough, Kaylee Farr, Marta Fay, Heidi Febinger, Jordan Finnell, Eric Fish, Henry Flores, Elyse Freeman, Briana Freshner, Carmen Frias, Andrew Galgoci, Alison Gardner, Jann Gardner, Michael Genin, Alex Gosch, Max Green, Denton Greenfield, Susana Grigorian, Sarah Guilmain, Matthew Guthrie, Michael Hancock, Chandana Haque, Michaela Hatch, Nathan Hatfield, Kristopher Howard, Gary Hsu, Sarah Hugo, Eric Hurst, Chock Ip, Evelyn Jaime, Jamie Jarvis, Christopher Johnson, Natalie Kalyuzhny, Tatjana Kanashiro, Ayla Khan, Matthew Kinn, Heather Kirkby, David Klitzing, Dylan Knutson, Eric Labonte, Dirk Lamb, Keirsten Lampkin, Tina Larson, Kevin Leggat, Jacob Letcher, Rebecca Levin, Mika Lindvall, Makail Lunt, Benjamin Mabey, Daniel Maljovec, Travis Martin, Katherine Matsumoto, Catherine Maxwell, Tarl McCallson, Ryan McComb, Britt McPartland, Jordan Mechanic, John Merrill, Ben Miller, Nathan Mitchell, Sara Moore, William Morrison, Kristen Morse, Khanhly Nguyen, Brandon Nichols, Scott Nielsen, Chad Nielsen, Lina Nilsson, Aaron Paugh, Kelsey Payne, Matthew Pfeiffer, Julia Pili, Kelly Porter, Bridget Pulver, Brandon Quach, Mark Rabe, Premraj Rajkumar, Alexis Ramos, Shelby Rampton, Brittnee Rhynes, Rosann Robinson, Juan Rodriguez Vera, Katrina Rodzon, Shane Rowley, Lourdes Rueda, Travis Rush, Kristen Rushton, Marissa Saunders, Collin Schlager, Spencer Schreier, Michael Secora, Michelle Sharer, Marika Xydes-Smith, Mark Smith, Rafael Soares, Ronald Spangle, Benjamin Sukow, Thos Swallow, James Taylor, Jordan Toutounchian, Katy Van Pelt, Luis Viramontez, Shafique Virani, Rebecca Webb, Nathan Wilkinson, Anthus Williams, Jathine Wong, Ann Marie Woodland, and Sijia Zhang, who have been working under extraordinary circumstances, home-schooling, socially-distancing and yet advancing this platform with incredible momentum. Further, we would like to acknowledge contributors at our Partner biosafety-level-3 facilities including Ryden Crowther, Ashley Dagley, Brett Hurst, Kie Hoon-Jung, John Morrey and Parker Weber at Utah State University and Alyssa Applegate, Richard Robison and Rebecca Scholl from Brigham-Young University.

## Author contributions

H.B.G., F.L.W., V.R., E.D.P., B.M.M, P.J. and N.D.C. designed and performed experiments. B.F., P.F.M., K.R.S., I.K.Q., N.H.L., S.W.G., J.F and J.D.J. performed computational analysis. C.A.B. and J.C. directed medicinal chemistry. K.R.T. and A.E.C advised and wrote. R.W.A., M.F.C., B.A.E., K.H. and conceived and designed experiments. M.A.S. conceived, designed and supervised experiments. A.S.L and S.S.H supervised the projects. M.L.V., I.S.H., Y.T.C and C.C.G conceived, designed, supervised experiments and wrote.

## SUPPLEMENTARY INFORMATION

**Fig. S1:**
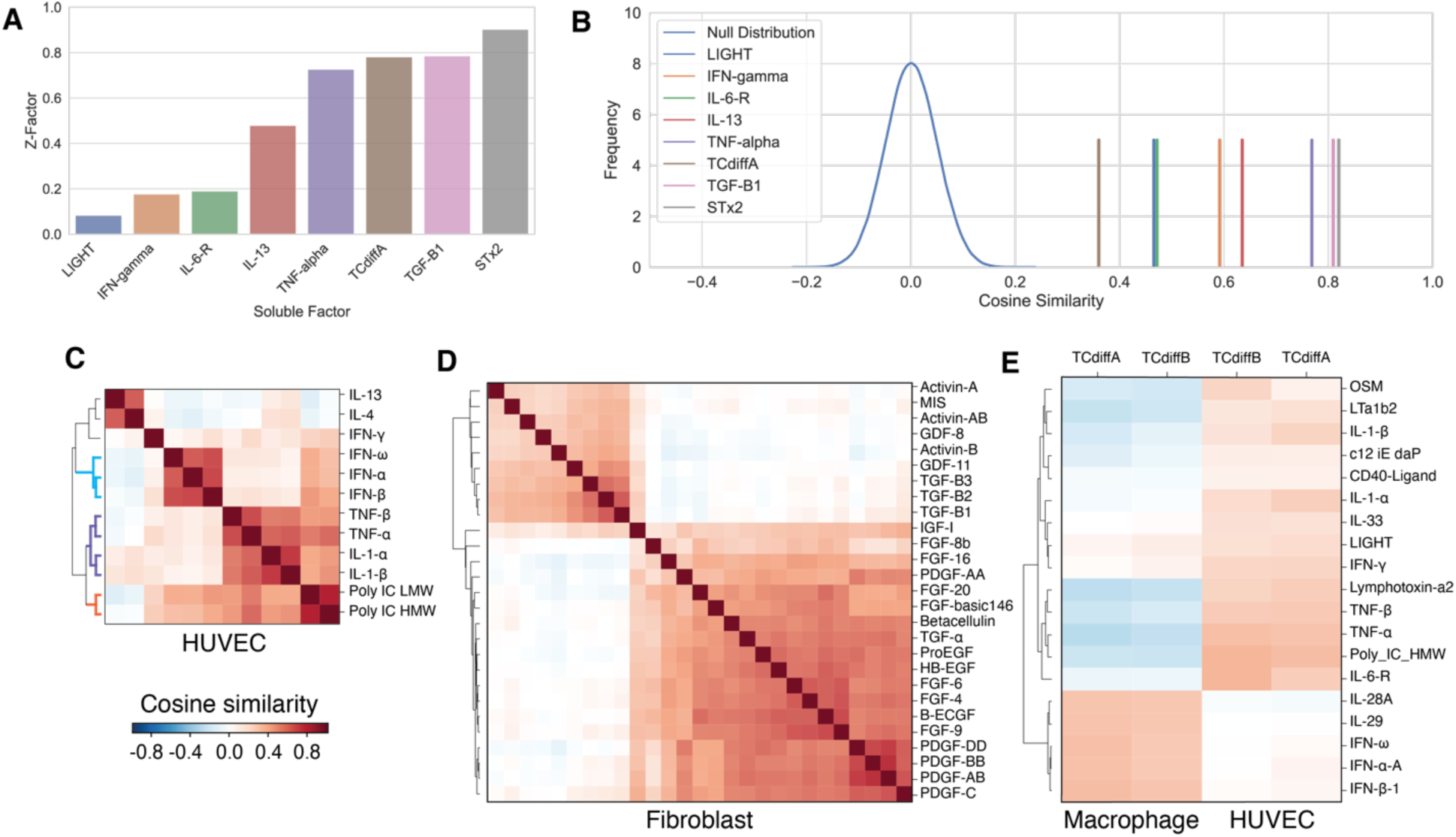
Determination of soluble factor phenoprints. **A**. Z-factor for the statistical confidence separating a series of soluble factors from untreated cells. **B**. For soluble factors presented in A, cosine similarity to an untreated null distribution, indicating strong separation for selected stimulant phenoprints. **C-E**. Hierarchical clustering of similarities. **c**. Phenoprints in HUVEC treated with three classes of factors: Interferons (blue segments), TLR3 agonists (orange segments), and NFκB-inducing cytokines (purple segments). **D**. Phenoprints in fibroblasts treated with growth factors. **e**. Differential response of *C. diff*. toxins A and B in HUVECs vs. macrophages.

**Fig. S2:**
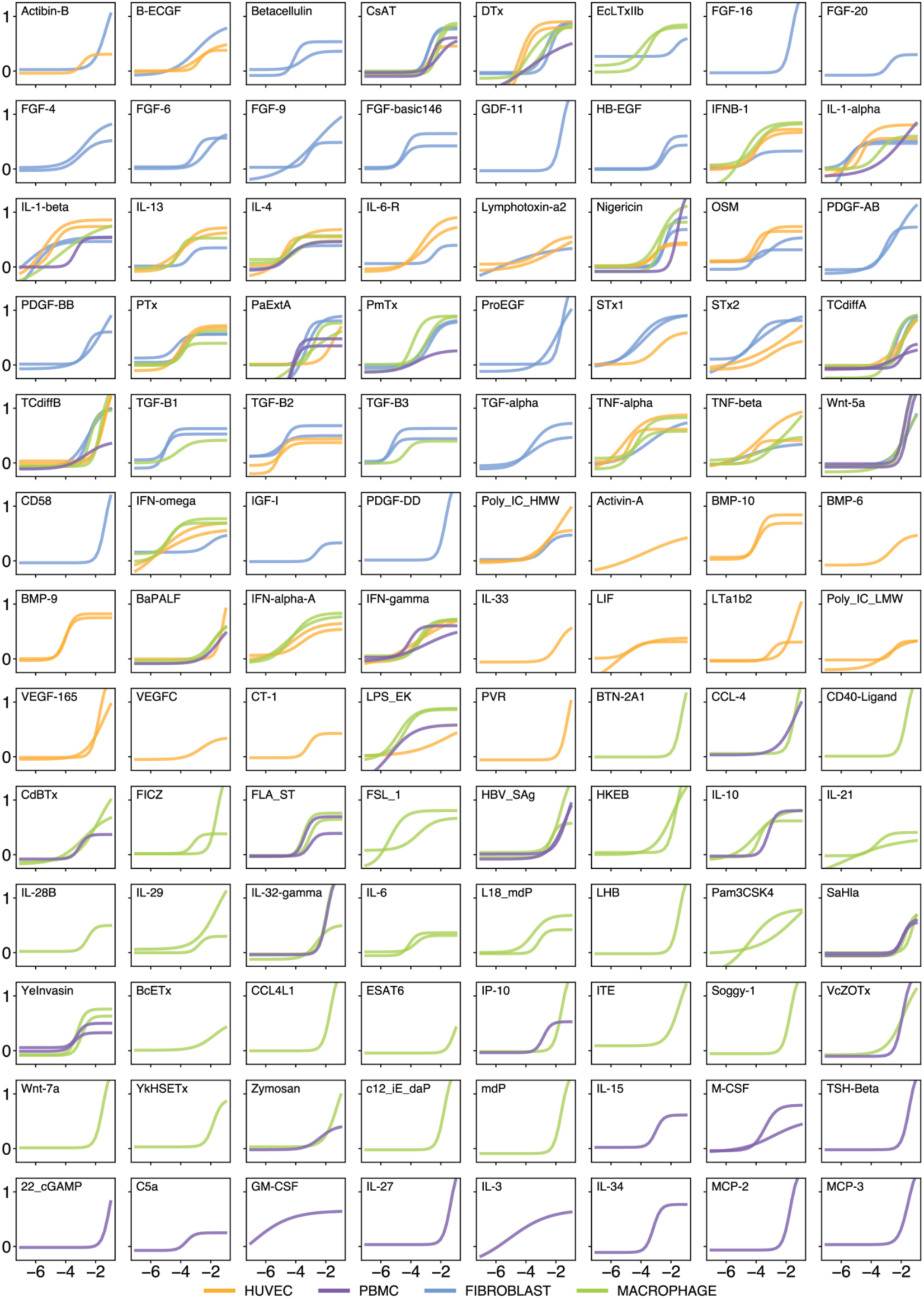
Cytokine Phenoprint Dose Curves. Convergence of phenoprints by dose, as represented by cosine similarity (y) of each dose (x, log10 concentration) to the highest dose for each reagent. Data is drawn from the union of two replicate experiments in each of 4 cell types (total 8 experiments): HUVEC, PBMC, Fibroblast, and Macrophage. The subset of fitted sigmoid curves for illustration were selected by filtering on a similarity threshold at any dose (cosine similarity >= 0.25) and fitted parameters (−7 < midpoint < −1.05; slope > 0.5; ceiling > 0.25; scale > 0.3). The 104 compounds shown represent 24% of the screened library.

**Fig. S3:**
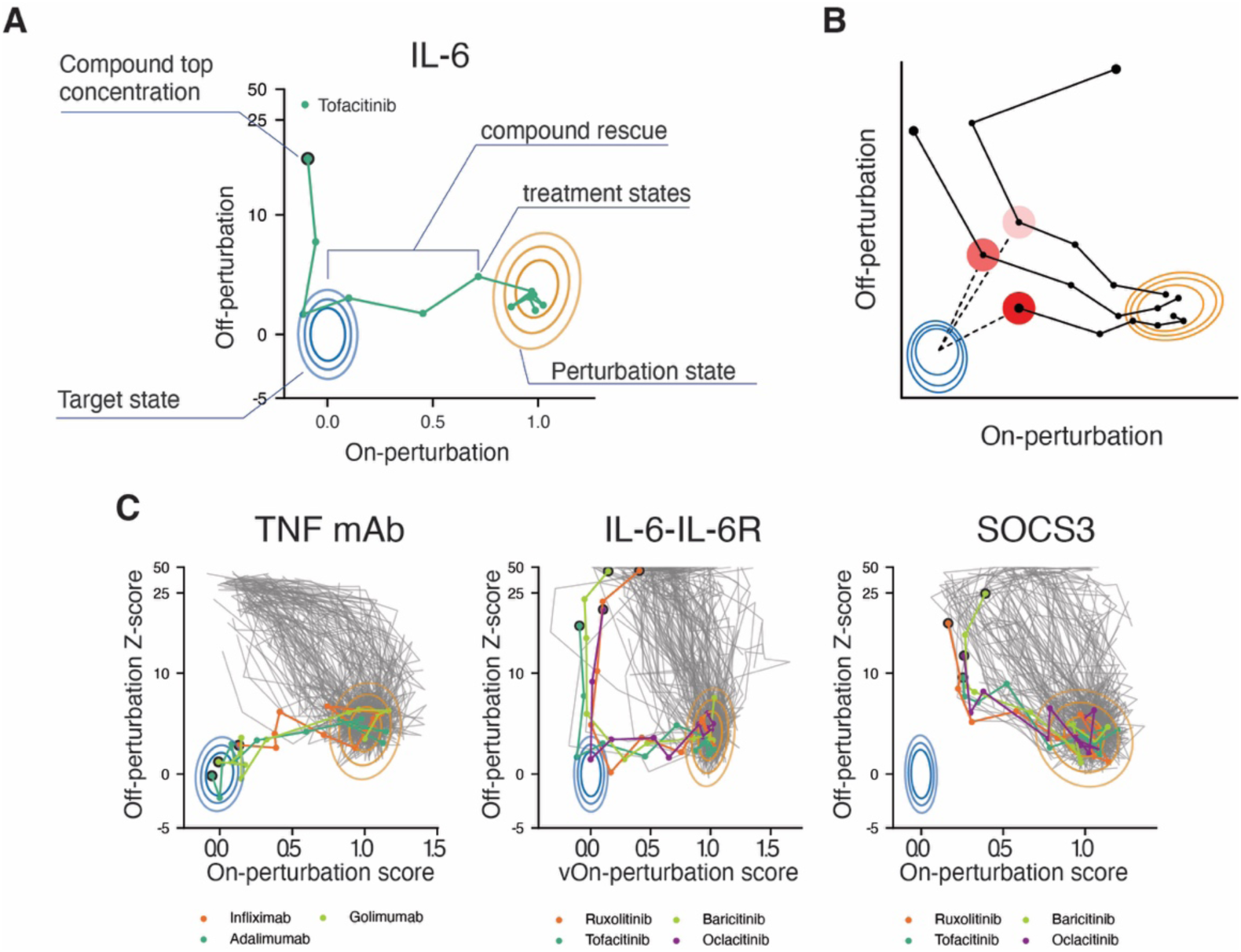
Projection in two dimensions and benchmark compound response. **A**. Schematic for interpreting projection of drug response in 2-dimensional plot. Contours show the 99%, 90%, and 50% distributions of on- and off-perturbation scores for the perturbation (orange) and target (blue) states. Ideal rescues are compounds that rescue along the on-perturbation x-axis toward the target state with minimal increase to the off-perturbation score on the y-axis. **B**. Potential hits are prioritized by the proximity of any dose to the target state, illustrated here with dashed lines and increasing red highlight intensity for higher-ranked dose-curve trajectories. **c**. Projections of treatments along the on-perturbation vector: rescue of the TNF-α phenoprint with clinically approved monoclonal antibodies, reversal IL-6-IL-6R receptor chimera, and reversal of SOCS3 CRISPR gene knockout. EC_50_ for high-dimensional compound rescue are indicated in parentheses

**Fig. S4:**
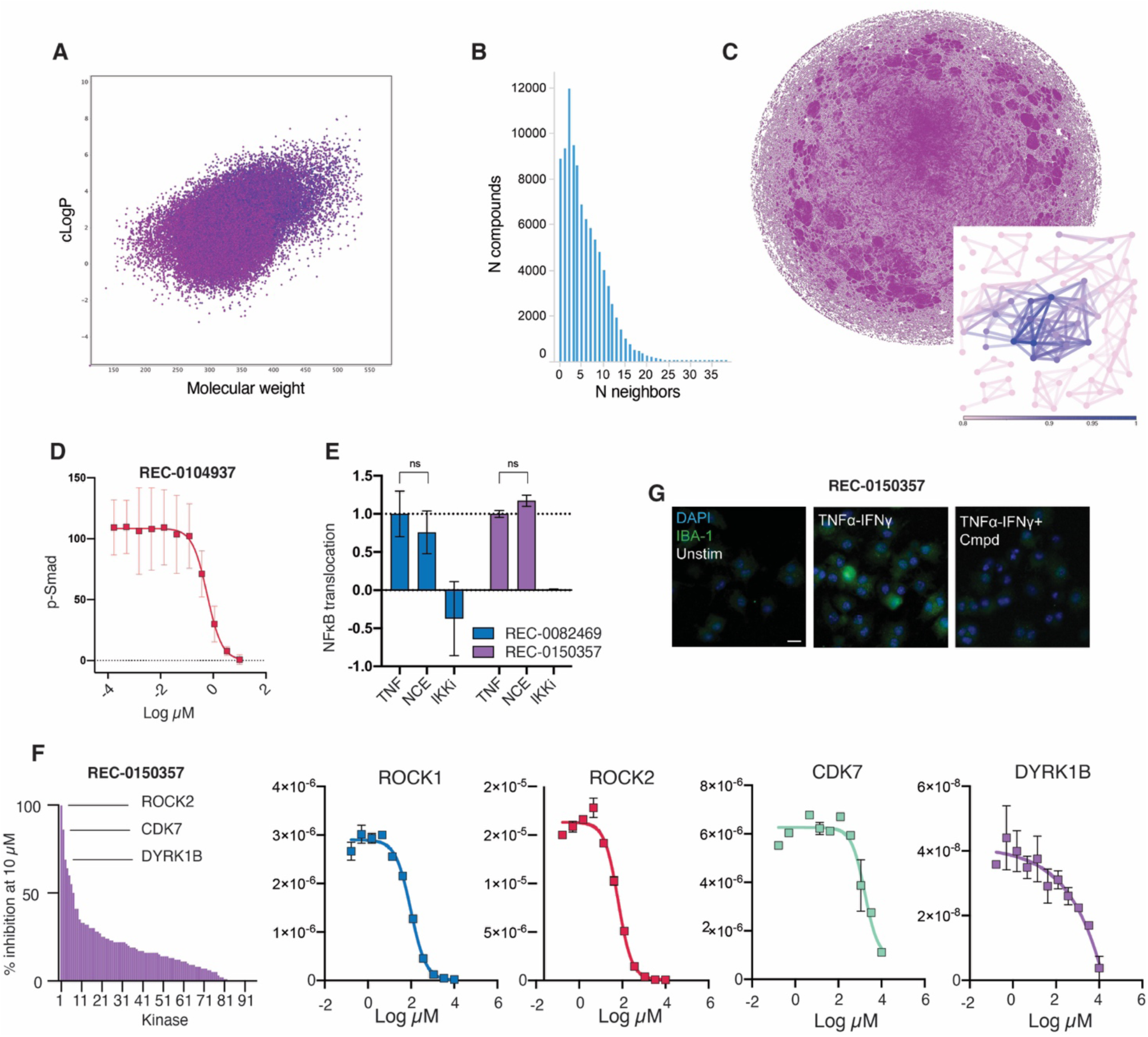
Library diversity and NCE compound validation. **A**. Library projected as molecular weight (x) and cLogP (y). **B**. Histogram of number of members of a structural scaffold. **C**. Pairwise assessment of scaffold diversity. Edges are projected between compounds >82% similar. Blue edges in the zoomed inset illustrates similarity to an example compound. **D**. Measurement of Smad phosphorylation in response to REC-0104937 in the presence of 1 ng/mL TGF-β-1. Microglial activation quantified by measurement of IBA-1 production. **E-F**. Kinase profiling of REC-0150357 in a single concentration screen (**E**) and concentration response (**F**). **G**. Representative images resulting from IBA-1 fluorescence immunostaining.

**Fig. S5:**
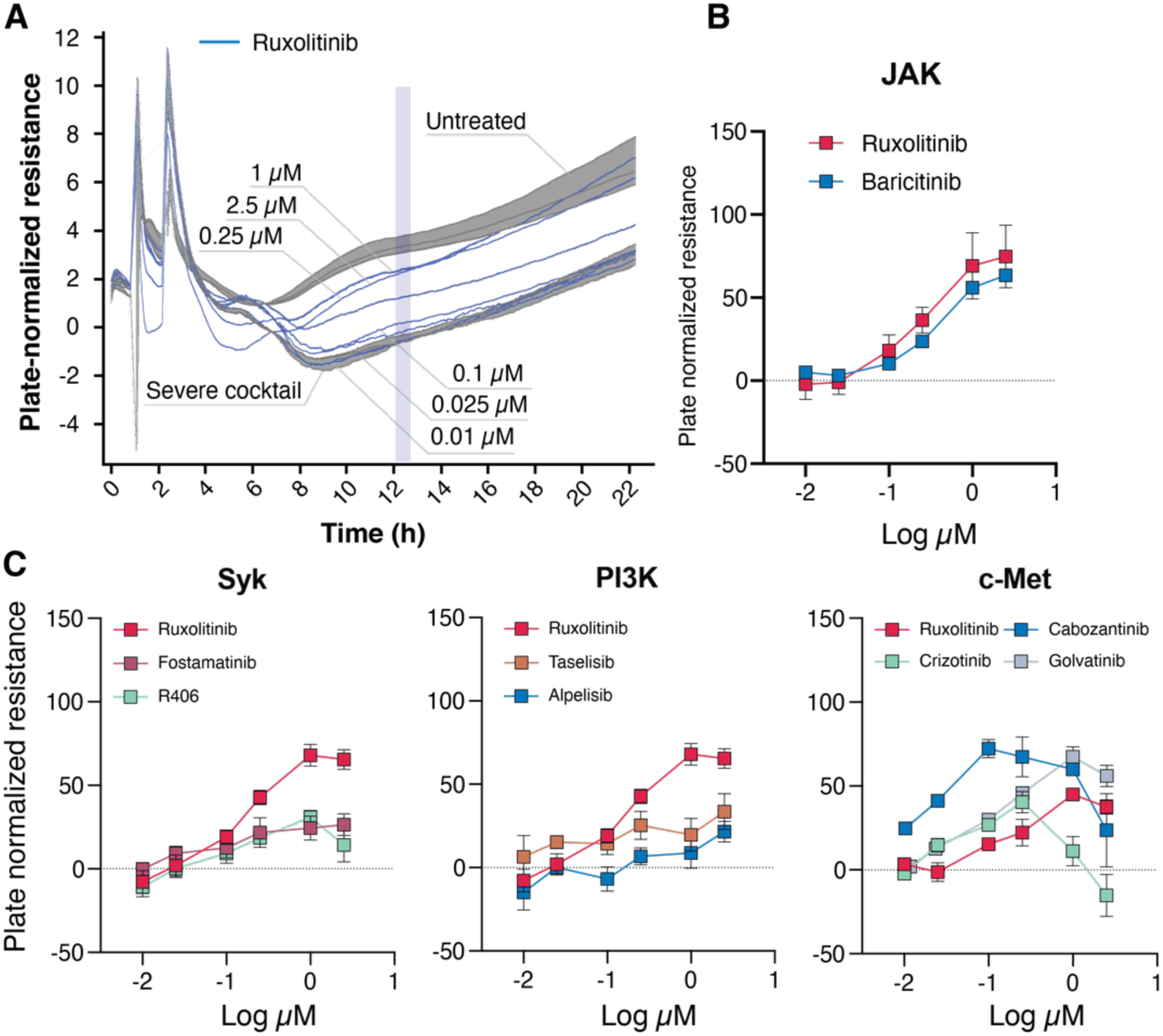
COVID-19-associated cytokine storm. **A**. ECIS trace for untreated and severe cocktail-treated wells. Blue lines represent concentrations of ruxolitinib. **B, C**. Protection of endothelial barrier integrity with active compounds. Data were averaged over a 12-minute window at hour 12 of ECIS measurement to visualize concentration response curves for the indicated compounds (n=5).

**Fig. S6:**
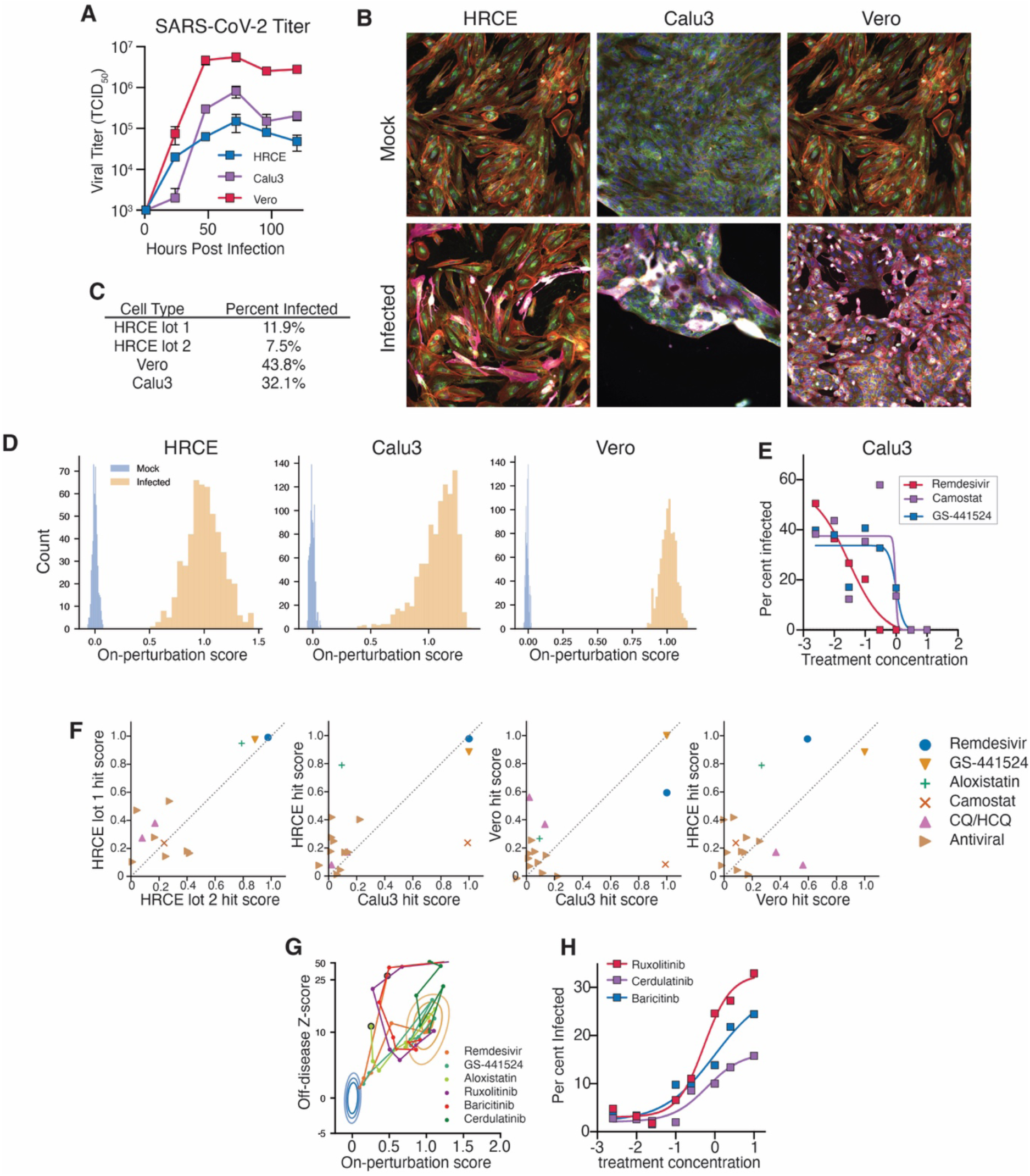
SARS-CoV-2 infection model. **A**. Quantification of active SARS-CoV-2 production over time in the indicated cell types using TCID50 measurement on Vero cells (n=2). **B**. Representative images of HRCE, Calu3 and Vero cells immunostained with SARS-CoV-2 nucleocapsid protein (pink) and modified cell paint dyes **C**. Infection rates of each tested cell type as analyzed by nucleocapsid immunostaining. Of note, HRCE donors displayed significant variation in infectability and only a minority of donors exhibited infection rates high enough for screening. Antibody stains were performed after the principal analysis concluded and are therefore not represented in the primary dataset used for phenoprint evaluation and compound screening. **D**. Infection of HRCE yielded a phenoprint against the mock-infected target population with an assay z-factor of 0.43 and was selected for further investigation. Vero and Calu3 cells also demonstrated screenable phenoprints. **E**. Quantification of percentage of cells infected using nucleocapsid protein immunostaining in Calu3 cells at 96 hours post infection for key compounds **F**. Consistency of hit scores for selected compounds across HRCE donors and between cell types. **G**. Projections of compound response of JAK inhibitor and control compounds onto the perturbation vector generated in SARS-CoV-2-infected HRCE. **H**. Quantification of percent of cells infected using nucleocapsid protein immunostaining in HRCE cells at 96 hours post infection for JAK inhibitors

**Table S1: Immune perturbants**

Names, catalogs and vendors for tested immune perturbants

**Table S2: Cellular phenotypes and** classes

Table to generate Fig. 2B

**Table S3: Cytokine storm cocktails**

Concentrations of individual factors in Cocktails representing circulating soluble factor levels for severe-COVID19 and healthy subjects.

